# Injury suppresses Ras cell competitive advantage through enhanced wild-type cell proliferation

**DOI:** 10.1101/2022.01.05.475078

**Authors:** Sara Gallini, Nur-Taz Rahman, Karl Annusver, David G. Gonzalez, Sangwon Yun, Catherine Matte-Martone, Tianchi Xin, Elizabeth Lathrop, Kathleen C. Suozzi, Maria Kasper, Valentina Greco

## Abstract

Healthy skin is a tapestry of wild-type and mutant clones. Although injury can cooperate with Ras mutations to promote tumorigenesis, the consequences in genetically mosaic skin are unknown. Here, we show that wild-type cells prevent oncogenic Ras-induced aberrant growth after injury. Although Hras^G12V/+^ and Kras^G12D/+^ cells outcompete wild-type cells in uninjured, mosaic tissue, their competitive advantage is suppressed after injury due to a selective increase in wild-type cell proliferation. EGFR inhibition abolishes the competitive advantage of wild-type cells after injury of Hras^G12V/+^-mosaic skin. Global loss of the cell cycle inhibitor p21 increases wild-type cell proliferation even without injury, suppressing the competitive advantage of Hras^G12V/+^ cells. Thus, injury plays an unanticipated role in switching the competitive balance between oncogenic and wild-type cells in genetically mosaic skin.

**One sentence Summary:** Injury-repair selectively induces wild-type cell proliferation to suppress oncogenic growth in Ras-mosaic skin epithelium.

## Main Text

Throughout our lifetimes, we acquire mutations in our skin, due to its constant exposure to environmental insults. As a result, phenotypically normal skin contains a mosaic of epithelial stem cells with somatic mutations, including in genes that are associated with cancer development such as the GTPase Ras family(*1, 2*). Constitutive activation of Ras oncogenes has been identified as the initial genetic event in 3-30% of human cutaneous squamous cell carcinomas (cSCCs)(*3-6*) and in experimentally induced cSCCs in mice(*7, 8*). In mouse models with mosaic epithelial expression of the constitutive active form of Hras (Hras^G12V/+^), mutant cells outcompete wild-type cells and expand in the uninjured skin epidermis(*9-11*). Although activated-Hras mutant cells are tolerated within otherwise wild-type and uninjured skin epithelium(*9-11*), injury has been shown to cooperate with oncogenic mutations to trigger tumorigenesis in various mouse models(*12-21*). We considered that the expansion of Hras^G12V/+^ cells in the epidermis could represent a vulnerability upon injury; Hras^G12V/+^ cells could futher expand and lead to tumors. For instance, the hyperproliferative environment generated during injury-repair may further stimulate the proliferative behavior of mutant cells and break the tolerance of the tissue. Here, we investigated how injury affects the oncogenic potential of Hras^G12V/+^ within a genetically mosaic and phenotypically relevant context.

### Injury-induced aberrant Hras^G12V/+^ growth is suppressed in mosaic skin

The stratified skin epidermis is uniquely accessible to direct observation, which allows the visualization of aberrant growth emergence at single-cell resolution. The basal layer contains epidermal stem cells, which can self-renew to generate more basal cells or differentiate and delaminate upwards to replace outer, barrier-forming cells(*22, 23*) (**Fig. S1A**). We hypothesized that injury-repair would cooperate with constitutive activation of the Hras oncogene (Hras^G12V/+^) to promote tumorigenesis in phenotypically normal genetically mosaic skin. To test this hypothesis, we generated mice in which we could induce and follow populations of Hras^G12V/+^ mutant cells within wild-type epithelium (*see Materials and Methods*; K14CreER; LSL-Hras^G12V/+^; LSL-tdTomato; K14H2B-GFP). In these mice, tamoxifen treatment activates Cre in Keratin 14-positive basal stem cells and, in turn, induces the co-expression of 1) Hras^G12V/+^ from its endogenous promoter and 2) a cytoplasmic fluorescent tdTomato reporter. Moreover, these mice express Histone H2B-GFP in basal cells, which perdures throughout differentiation(*24*), allowing the visualization of all basal stem cells and their progeny (**Fig. 1A**). We treated mice with tamoxifen at 3 weeks of age and, three days later, introduced a full-thickness injury down to the cartilage (4 mm diameter punch biopsy) in one ear. We employed two doses of tamoxifen to drive Hras^G12V/+^ expression in either approximately 99% of basal stem cells (Hras^G12V/+^-max) to recapitulate previous studies of homogeneous models(*12-14, 25, 26*), or approximately 65% of basal stem cells (Hras^G12V/+^-mosaic) to mimic genetically mosaic skin (**Fig. 1B**). As a control, we also engineered K14CreER; LSL-tdTomato; K14H2B-GFP mice and treated them similarly to drive tdTomato expression in approximately 65% of wild-type basal stem cells (wild-type-mosaic) (**Fig. 1B**).

**Fig. 1:**
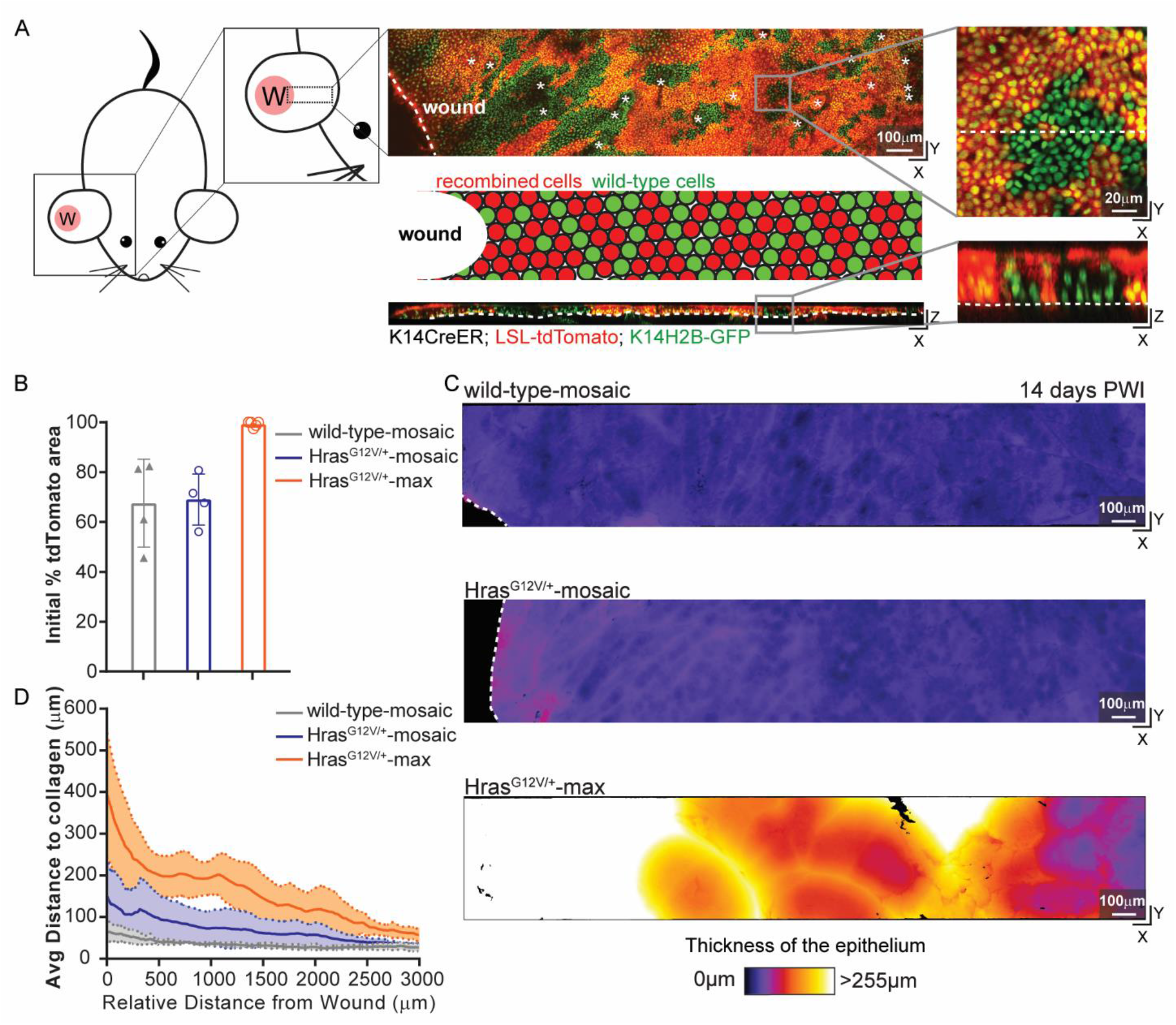
Injury-induced aberrant Hras^G12V/+^ growth is suppressed in mosaic skin. **A)** *Left*: cartoon schematic depicting a 4 mm punch biopsy, full-thickness wound, on one mouse ear and the area around the wound imaged by two-photon microscopy. *Right:* top down (*x-y*) and transversal (*x-z*) views of a two-photon image of the skin epithelium at 14 days post-wound induction (PWI) of K14CreER; LSL-tdTomato; K14H2B-GFP mouse (asterisks mark hair canals and dashed lines mark the basement membrane in *x-z* view and the wound edge in *x-y* view; Scale bar=100 µm) and (*middle*) a cartoon schematic of wild-type (green) and recombined cells post-tamoxifen injection (red) around the injury. Tertiary inset: close-up images of top down (*x-y*) and transversal (*x-z*) views of the skin epithelium show all the epithelial cell nuclei (K14H2B-GFP) in green and recombined cells, expressing tdTomato, in red (dashed lines mark the transversal section in *x-y* view and the basement membrane in *x-z* view; Scale bar=20 µm). **B)** Initial percentage of tdTomato+ area in the first revisit at 3 days PWI of wild-type-mosaic, Hras^G12V/+^-mosaic and Hras^G12V/+^-max (n=4 wild-type-mosaic and Hras^G12V/+^-mosaic mice and n=5 Hras^G12V/+^-max mice). **C)** Heat maps of the top down (*x-y*) view of representative two-photon images around the injury in wild-type-mosaic, Hras^G12V/+^-mosaic and Hras^G12V/+^-max at 14 days PWI (dashed lines highlight the wound edge). The color code corresponds to epithelial thickness and is used to identify the presence of aberrant growth around the injury (see key below images). Only Hras^G12V/+^-max developed aberrant oncogenic growth around the injury. Scale bar=100 µm. **D)** Quantification of the average thickness, using IMARIS and MatLab software, of the epithelium at 14 days PWI at different distances from the edge of the wound in wild-type-mosaic, Hras^G12V/+^-mosaic and Hras^G12V/+^-max. Solid lines represent means and dashed lines standard deviations. n=4 wild-type-mosaic and Hras^G12V/+^-mosaic mice and n=5 Hras^G12V/+^-max mice.

We monitored the injured epithelium over time by combining deep tissue imaging with quantitative analyses via IMARIS and MatLab software, which allowed us to evaluate tissue thickness with intensity heat maps (**Fig. 1C, D, Movie S1, 2, 3, 4**, *see Materials and Methods*). Intriguingly, at 14 days post-wound induction (PWI), the Hras^G12V/+^-mosaic models did not develop the aberrant growth and thick epithelium observed in Hras^G12V/+^-max models (**Fig. 1C, D, Movie S1, 2, 3, 4**). Histopathological analysis further showed that at day-14 PWI, the skin epithelium around the repaired injury in the Hras^G12V/+^-mosaic model was normal, despite the high burden of Hras^G12V/+^ mutation (**Fig. S1B, C, E**). In contrast, abnormal growth formed rapidly within the two weeks after injury induction in Hras^G12V/+^-max, as expected (**Fig. 1C, D, Fig. S1D, E, Movie S3, 4**). Collectively, these data show that Hras^G12V/+^ cells break homeostatic tissue architecture during injury-repair only when nearly all the basal stem cells express Hras^G12V/+^.

### Injury-repair alters the competitive balance between wild-type and Hras^G12V/+^ cells in mosaic skin

Having discovered that injury-repair does not trigger aberrant growth in Hras^G12V/+^-mosaic tissue, we next investigated how Hras^G12V/+^ and wild-type cells within mosaic epithelia respond to injury. Our previous study showed that embryonically induced Hras^G12V/+^ basal stem cells integrate and expand in the skin epidermis, eventually outcompeting wild-type cells(*9*). We had hypothesized that this proliferative advantage of Hras^G12V/+^ cells would be amplified during injury-repair, which has a higher proliferative demand than uninjured skin. To test our hypothesis, we revisited the same wild-type and Hras^G12V/+^ cells in the skin epidermis of live mice for one month, with or without injury-repair (**Fig. 2A, Fig. S2A**). Epithelial cells start to contribute to re-epithelialization approximately three days after injury(*27, 28*). Therefore, we started our analysis three days PWI (6 days after tamoxifen-induced mosaicism, **Fig. 2A**). We drew boundaries between GFP+/tdTomato+ and GFP+/tdTomato− regions and represented the tdTomato+ areas as a percentage of the total area quantified (**Fig. 2B, C, D, E**).

**Fig. 2:**
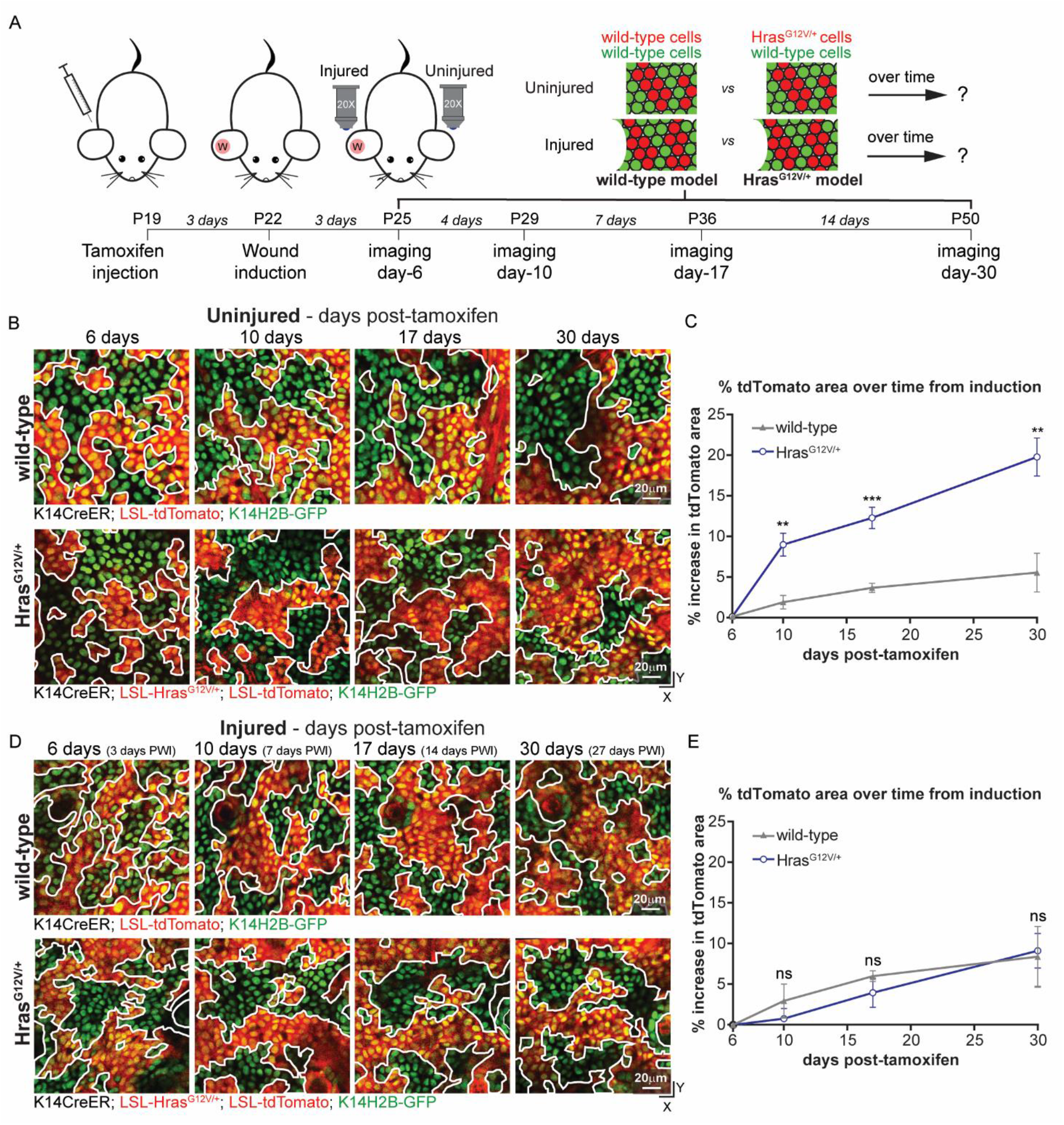
Injury-repair alters the competitive balance between wild-type and Hras^G12V/+^ cells in mosaic skin. All epithelial nuclei are in green (K14H2B-GFP) and recombined cells are in red (LSL-tdTomato). **A)** Schematic representation of the experimental design: induction of recombination through tamoxifen injection at post-natal day 19, followed by injury induction 3 days later and imaging of uninjured and injured areas at 6 days, 10 days, 17 days and 30 days post-recombination. GFP+/tdTomato+ (red) and GFP+/tdTomato− (green) cells in wild-type and Hras^G12V/+^-mosaic are tracked and compared over time. **B)** Two-photon representative revisit images of the same area of the basal stem cell layer of the epidermis in uninjured condition in wild-type and Hras^G12V/+^-mosaic at 6 days, 10 days, 17 days and 30 days post-tamoxifen injection. Hras^G12V/+^ cells (GFP+/tdTomato+) outcompeted wild-type neighbors (GFP+/tdTomato−) over time. White lines highlight the boundaries between GFP+/tdTomato+ and GFP+/tdTomato− populations. **C)** Percentage increase of tdTomato+ area over time in uninjured condition in wild-type and Hras^G12V/+^-mosaic. Initial time point is at 6 days post-tamoxifen injection followed by 10 days, 17 days and 30 days from the recombination (*see Materials and Methods*). **D)** Two-photon representative revisit images of the same area of the basal stem cell layer of the epidermis during injury-repair in wild-type and Hras^G12V/+^-mosaic at 3 days, 7 days, 14 days and 27 days PWI. In injured condition, Hras^G12V/+^ cells (GFP+/tdTomato+) did not outcompete wild-type neighbors (GFP+/tdTomato−) over time. **E)** Same quantification as in C) at 3 days, 7 days, 14 days and 27 days PWI. Statistics: unpaired, two-tailed *t-test* between wild-type and mutant mice at different time points in uninjured and injured conditions. **, P<0.005 and ***, P<0.0005, ns indicates not statistically significant. Data are represented as means and standard deviations. n=4 mice for each condition. Scale bar=20 µm.

We found that the Hras^G12V/+^/tdTomato+ population expanded substantially in uninjured Hras^G12V/+^-mosaic epithelium, much more than the tdTomato+ population in uninjured wild-type-mosaic epithelium. After one month, Hras^G12V/+^ cells outcompeted wild-type cells and increased their occupancy of the basal stem cell layer by approximately 20% in uninjured Hras^G12V/+^-mosaic mice, consistent with our previous work(*9*)(**Fig. 2B, C, Fig. S2B**). In contrast, one month after injury, the Hras^G12V/+^/tdTomato+ and wild-type/tdTomato+ percentages were similar. This finding was unexpected and indicates that Hras^G12V/+^ cells failed to outcompete wild-type cells and expand after injury-repair of Hras^G12V/+^-mosaic mice (**Fig. 2D, E, Fig. 1B**). Collectively, these data demonstrate that the injury-repair process does not amplify but rather abrogates the competitive advantage that Hras^G12V/+^ cells have over wild-type cells in the absence of injury.

### Injury selectively induces the proliferation of wild-type cells in Hras^G12V/+^-mosaic skin

The suppressed expansion of Hras^G12V/+^ cells after injury prompted us to investigate how this process affects different cellular behaviors of mutant and wild-type cells. Specifically, we first examined proliferation, given that Ras is a key regulator of epithelial cell proliferation in the skin epithelium. Indeed, epithelial stem cells *in vitro* and *in vivo* fail to proliferate upon ablation of all Ras isoforms(*29*). To examine the proliferation rate, we scored mitotic cells in uninjured or injured skin by immunostaining for the mitotic marker phospho-Histone-3. We observed an increase in epithelial cell proliferation accompanying efficient wound repair at 3-days PWI in the wild-type-mosaic model (**Fig. 3A, B**). In sharp contrast, although we observed an increase in the mitotic events of wild-type cells in the Hras^G12V/+^-mosaic model, Hras^G12V/+^ cell proliferation was unaltered during repair (**Fig. 3B**). These findings were corroborated by measuring mitotic figures (**Fig. S3A, B, C**). Therefore, wild-type cells have an unexpected and selective proliferative advantage over Hras^G12V/+^ cells in the acute phase of injury-repair.

**Fig. 3:**
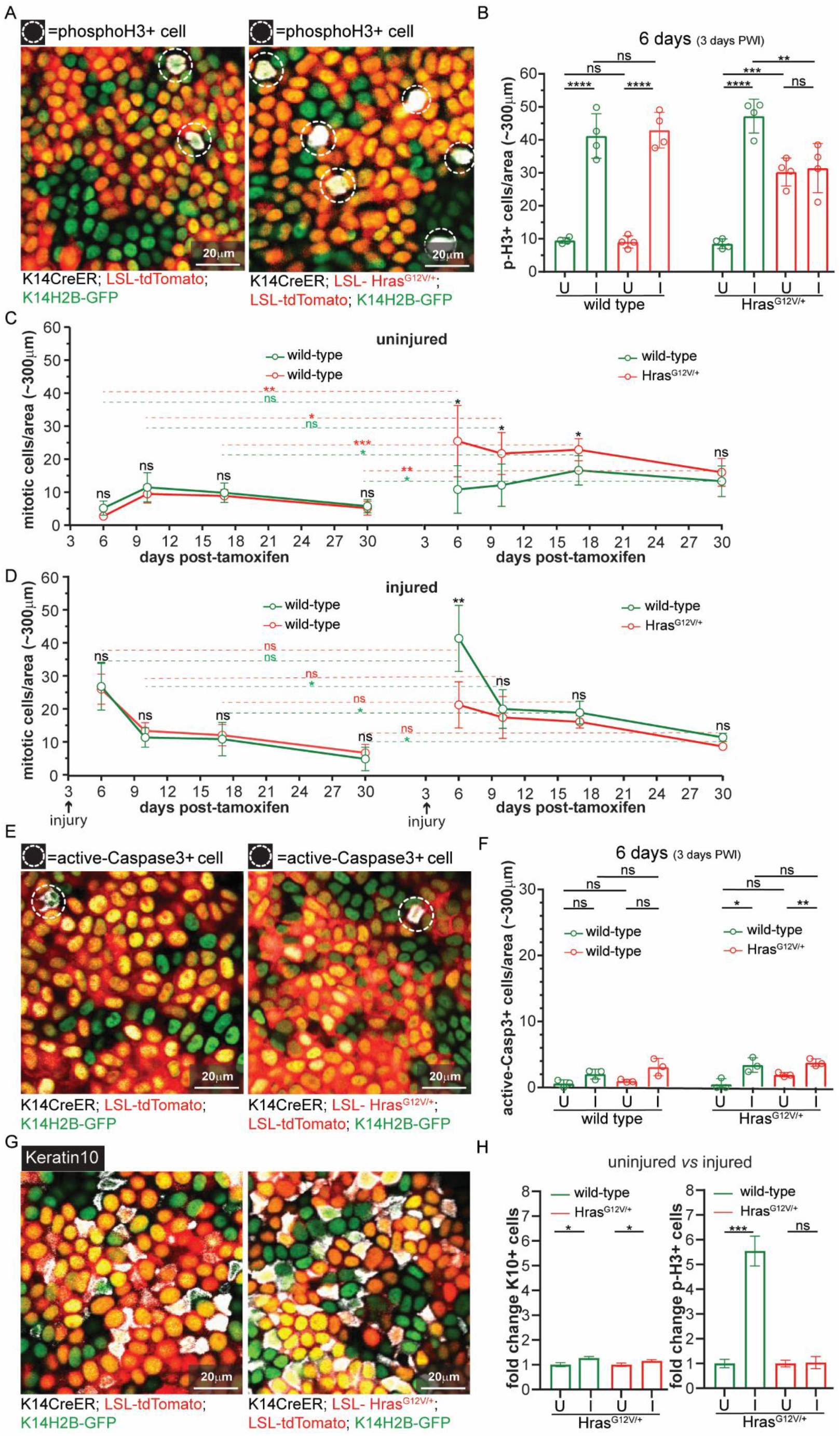
Injury selectively induces the proliferation of wild-type cells in Hras^G12V/+^-mosaic skin. All epithelial nuclei are in green (K14H2B-GFP) and recombined cells are in red (LSL-tdTomato). **A)** Two-photon representative images of the epidermal preparation staining against phospho-Histone-3, a mitotic marker, in both wild-type and Hras^G12V/+^-mosaic (positive cells highlighted with white dashed circles). **B)** Quantification of phospho-Histone-3 positive cells in GFP+/tdTomato+ and GFP+/tdTomato− populations in both wild-type and Hras^G12V/+^-mosaic at 6 days after tamoxifen treatment in injured/3 days PWI (I) and uninjured (U) ears. The number of positive cells in the immunostaining was divided by the GFP+/tdTomato+ and GFP+/tdTomato− areas and multiplied by the total area analyzed (∼300 µm^2^) to compare the two populations. We analyzed three ∼300 µm^2^ areas for each mouse. n=4 mice for each condition. **C)** Quantification of mitotic figures in uninjured ears at 6 days, 10 days, 17 days and 30 days post-tamoxifen injection in GFP+/tdTomato+ and GFP+/tdTomato− populations in wild-type-mosaic (*left*) and Hras^G12V/+^-mosaic (*right*). Statistical analysis of mitotic events between wild-type-mosaic and Hras^G12V/+^-mosaic in GFP+/tdTomato− (green) or GFP+/tdTomato+ (red) population over time are represented with green or red dotted lines. n=4 mice for each condition. **D)** Same quantification as in C) in injured ears. **E)** Two-photon representative images of the epidermal preparation staining against active-Caspase-3, an apoptotic marker, in both wild-type and Hras^G12V/+^-mosaic (positive cells highlighted with a white dashed circle). **F)** Quantification of active-Caspase-3 positive cells in GFP+/tdTomato+ and GFP+/tdTomato− areas in both wild-type and Hras^G12V/+^-mosaic in injured/3 days PWI (I) and uninjured (U) ears. The number of positive cells in the immunostaining was divided by the GFP+/tdTomato+ and GFP+/tdTomato− areas and multiplied by the total area analyzed (∼300 µm^2^) to compare the two populations. We analyzed three ∼300 µm^2^ areas for each mouse. n=3 mice for each condition. **G)** Two-photon representative images of the epidermal preparation staining against Keratin10, an early differentiation marker, in both wild-type and Hras^G12V/+^ -mosaic (positive cells are stained in white). **H)** Quantification of the fold change differences in the number of cells expressing Keratin10 (*left*) or phospho-Histone3 (*right*) between uninjured and injured conditions in GFP+/tdTomato+ and GFP+/tdTomato− populations in Hras^G12V/+^-mosaic. Uninjured condition is equal to 1. n=3 mice for Keratin10 staining in each condition. n=4 for phospho-Histone-3 staining in each condition. Statistics: Pair, two-tailed *t-test*, for comparison between GFP+/tdTomato+ and GFP+/tdTomato− populations in the same group of mice. Unpaired, two-tailed *t-test* for comparison between GFP+/tdTomato+ and GFP+/tdTomato− populations in different groups of mice. *, P<0.05, **, P<0.005 and ***, P<0.0005 and ****, P<0.00001. ns indicates not statistically significant. Data are represented as means and standard deviations. Scale bar=20 µm.

To determine if the increased proliferation of wild-type cells that we observed at 3 days PWI was sustained over time, we scored mitotic events for over four weeks. In the wild-type-mosaic model, the initial increase in proliferation observed in both tdTomato+ and tdTomato− wild-type cells returned to baseline by 7 days PWI (10 days after tamoxifen-induced mosaicism) and looked similar to uninjured mice as expected (**Fig. 3C, D)**. The initially elevated wild-type cell proliferation observed in the injured Hras^G12V/+^-mosaic model also decreased over time, and eventually looked similar to Hras^G12V/+^ neighbors, but still greater than wild-type cell proliferation after injury of the wild-type-mosaic model (**Fig. 3C, D**). In contrast, the proliferative capacity of Hras^G12V/+^ cells in the Hras^G12V/+^-mosaic model was not substantially affected by injury-repair at any of the time points analyzed (**Fig. 3C, D**). Thus, injury induced a persistent increase in wild-type cell proliferation in the Hras^G12V/+^-mosaic model but not in the wild-type-mosaic model (**Fig. 3D**). The balanced proliferation of wild-type and Hras^G12V/+^ cells sustained at later time points after injury would effectively continue to prevent the expansion of Hras^G12V/+^ cells in the Hras^G12V/+^-mosaic model.

To test whether injury-repair leads to genotype-specific changes in other cellular behaviors, we monitored apoptosis, which is an established cell competition mechanism(*30*) and inhibited by Ras signaling(*31, 32*). We examined cell death by scoring for either nuclear fragmentation events or expression of an apoptotic marker, active-Caspase-3. The overall frequency of apoptosis was low, and we did not observe significant differences in cell death events of wild-type or Hras^G12V/+^ cells in mice with or without injury at 6 days post-tamoxifen-induced mosaicism and at later time points (**Fig. 3E, F, Fig. S3D, E, F, G**). Differentiation is another mechanism of cell loss that could influence wild-type or Hras^G12V/+^ cell competition in the skin epidermis. To comprehensively evaluate differentiation rates, we interrogated the expression of early differentiation markers by both protein and scRNA-seq analyses (**Fig. 3G, H, Fig. S3H, Fig. S4, Fig. S5**). scRNA-seq analysis revealed an earlier onset of differentiation during injury-repair in the Hras^G12V/+^-mosaic model compared to the wild-type-mosaic model, inferred from an increase of differentiation markers and a decrease of stemness transcripts (*see Materials and Methods;* **Fig. S5C)**. To determine if the increased differentiation depended on the genotype, we quantified the expression of the early differentiation marker Keratin10 in wild-type and Hras^G12V/+^ cells. We observed a similar increase of differentiating cells upon injury for both wild-type and Hras^G12V/+^ cells compared to the uninjured condition, in contrast to the selective increase in proliferation of wild-type cells (**Fig. 3G, H, Fig. S3H**).

Collectively, these results show that injury-repair increases wild-type cell proliferation only, and therefore suppresses Hras^G12V/+^ cell expansion in the Hras^G12V/+^-mosaic model.

### Aggressive Kras^G12D/+^ mutant cells lose their competitive advantage during injury-repair of mosaic skin

Next, we investigated whether the injury-repair process in mosaic skin could effectively suppress a more aggressive allele of the Ras gene family, Kras^G12D^. Mice with homogeneous activation of Kras^G12D/+^ in the skin epidermis rapidly develop oncogenic growth in areas of constant abrasion (data not shown; (*33, 34*)). Kras is one of the most frequently mutated oncogenes in human cancer and broadly activated across epithelial cancers, including cSCCs(*35, 36*). We generated mice in which we could induce and follow Kras^G12D/+^ cells within wild-type epithelium (*see Materials and Methods*; K14CreER; LSL-Kras^G12D/+^; LSL-tdTomato; K14H2B-GFP). Similar to Hras^G12V/+^/tdTomato+ cells in the Hras^G12V/+^-mosaic model, Kras^G12D/+^/tdTomato+ cells expanded in uninjured Kras^G12D/+^-mosaic mouse skin but did not expand after injury (**Fig. 4A, B, Fig. S6A**). We scored mitotic events before and after injury induction, and again observed a selective increase in the proliferation of wild-type cells but not mutant cells in Kras^G12D/+^-mosaic mouse skin, similar to what we observed in the Hras^G12V/+^-mosaic model (**Fig. 4C**).

**Fig. 4:**
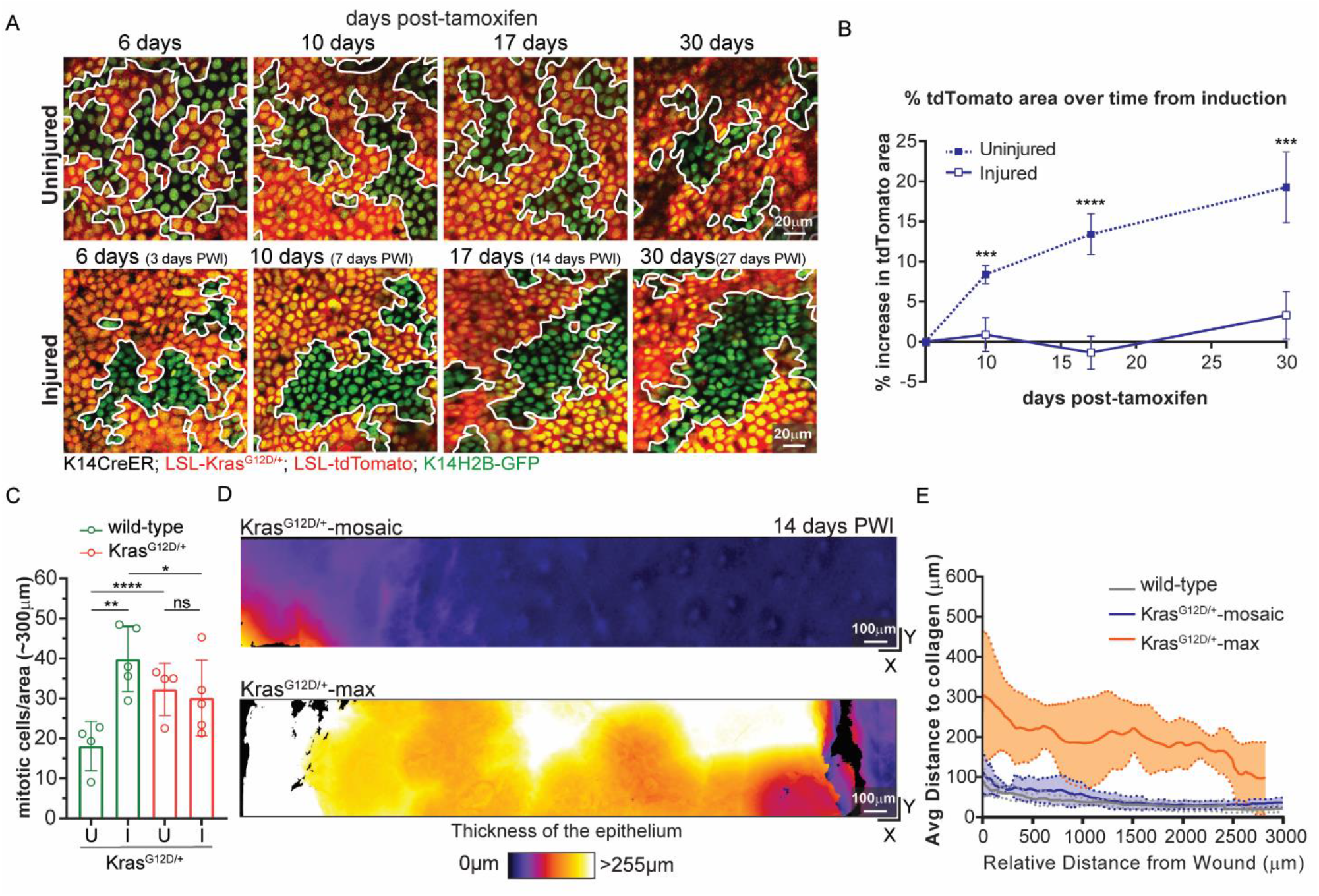
Aggressive Kras^G12D/+^ mutant cells lose their competitive advantage during injury-repair of mosaic skin. All epithelial nuclei are in green (K14H2B-GFP) and recombined cells are in red (LSL-tdTomato). **A)** Two-photon representative revisit images of the same area of the basal stem cell layer of the epidermis in Kras^G12D/+^-mosaic in uninjured (*top row*) and injured (*bottom row*) conditions. Kras^G12D/+^ cells (GFP+/tdTomato+) outcompeted wild-type neighbors (GFP+/tdTomato−) over time in uninjured condition but not during injury-repair. White lines highlight the boundaries between GFP+/tdTomato+ and GFP+/tdTomato− populations. Scale bar=20 µm. **B)** Percentage increase of tdTomato+ area at 6 days (3 days PWI), 10 days (7 days PWI), 17 days (14 days PWI) and 30 days (27 days PWI) post-tamoxifen injection in uninjured and injured conditions. n=4 uninjured Kras^G12D/+^-mosaic mice and n=5 injured Kras^G12D/+^-mosaic mice. **C)** Quantification of mitotic figures in GFP+/tdTomato+ and GFP+/tdTomato− areas in both wild-type and Kras^G12D/+^-mosaic. The number of mitotic events was divided by the GFP+/tdTomato+ and GFP+/tdTomato− areas and multiplied by the total area analyzed (∼300 µm^2^) to compare the two populations. We analyzed three ∼300 µm^2^ areas for each mouse. n=4 uninjured Kras^G12D/+^-mosaic mice and n=5 injured Kras^G12D/+^-mosaic mice. **D)** Heat maps of the top down (*x-y*) view of representative two-photon images around the injury in Kras^G12D/+^-mosaic and Kras^G12D/+^-max at 14 days PWI. The color code corresponds to epithelial thickness (see key below images) and is used to identify the presence of aberrant oncogenic growth around the injury. Only Kras^G12D/+^-max developed aberrant growth around the injury. **E**) Quantification of the average thickness, using IMARIS and MatLab software, of the epithelium at 14 days PWI at different distances from the edge of the wound in wild-type, Kras^G12D/+^-mosaic and Kras^G12D/+^-max. Solid lines represent means and dashed lines standard deviations. n=4 Kras^G12D/+^-mosaic mice and n=3 wild-type and Kras^G12D/+^-max mice. Statistics: Pair, two-tailed *t-test*, for comparison between GFP+/tdTomato+ and GFP+/tdTomato− populations in the same group of mice. Unpaired, two-tailed *t-test* for comparison between GFP+/tdTomato+ and GFP+/tdTomato− populations in different groups of mice and between Kras^G12D/+^-mutant mice in uninjured and injured conditions at different time points. *, P<0.05, **. P<0.005, ***, P<0.0005 and ****, P<0.00001, ns indicates not statistically significant. Data are represented as means and standard deviations.

To monitor phenotypes at the tissue level, we applied two-photon microscopy with quantitative analyses of epidermal thickness represented by intensity heat maps. Despite the high burden of Kras^G12D/+^ mutation (approximately 65% of recombined cells), the skin epithelium of Kras^G12D/+^-mosaic mice remained similar to wild-type-mosaic models after injury (**Fig. 1C, Fig. 4D, E, Fig. S6A, B**). In contrast, the Kras^G12D/+^-max model, in which nearly all the basal stem cells expressed Kras^G12D/+^, displayed rapid abnormal growth within the first two weeks after injury (**Fig. 4D, E, Fig. S6A, C, D**).

Overall, our work strongly suggests that the selective increase in wild-type cell proliferation during injury-repair of mosaic skin limits the expansion and tumorigenic potential of mutant cells expressing different oncogenic variants of the Ras gene family.

### EGFR inhibition promotes the expansion of Hras^G12V/+^ cells in mosaic skin after injury

We found that injury-repair in mosaic skin triggers a specific increase in the proliferation of wild-type cells but not Kras^G12D/+^ and Hras^G12V/+^ cells. However, it remained unclear whether the increased proliferation of wild-type cells *per se* suppressed the competitive advantage of Ras-mutant cells. To investigate this, we specifically reduced the proliferation of wild-type cells by inhibiting the epidermal growth factor receptor (EGFR). In wild-type cells, EGFR activation is required to upregulate Ras signaling and thus promote cell proliferation of the skin epidermis during injury-repair (*37-40*). In contrast, Hras^G12V/+^ cells are less dependent on EGFR activation for Ras signaling, due to their expression of constitutively active Ras^G12V^. Therefore, EGFR inhibition should selectively reduce proliferation of wild-type cells after injury. We employed the EGFR inhibitor Gefitinib, which we verified repressed EGFR activity during injury-repair of wild-type-mosaic mice (*see Materials and Methods*; **Fig. S7A**). As expected, Gefitinib treatment selectively inhibited the proliferation of wild-type cells, but not of Hras^G12V/+^ cells, in Hras^G12V/+^-mosaic models after injury (**Fig. 5A**). To assess how Gefitinib treatment affects cell competition during injury-repair, we tracked the percentage of surface coverage of Hras^G12V/+^ cells (GFP+/tdTomato+) at 3-, 7- and 14-days PWI (6-, 10-, 17-days post-tamoxifen-induced mosaicism). Strikingly, we discovered that EGFR inhibition reestablishes the competitive advantage of Hras^G12V/+^ cells over their wild-type neighbors during injury-repair of mosaic mice (**Fig. 5B, C, Fig. S7B**).

**Fig. 5:**
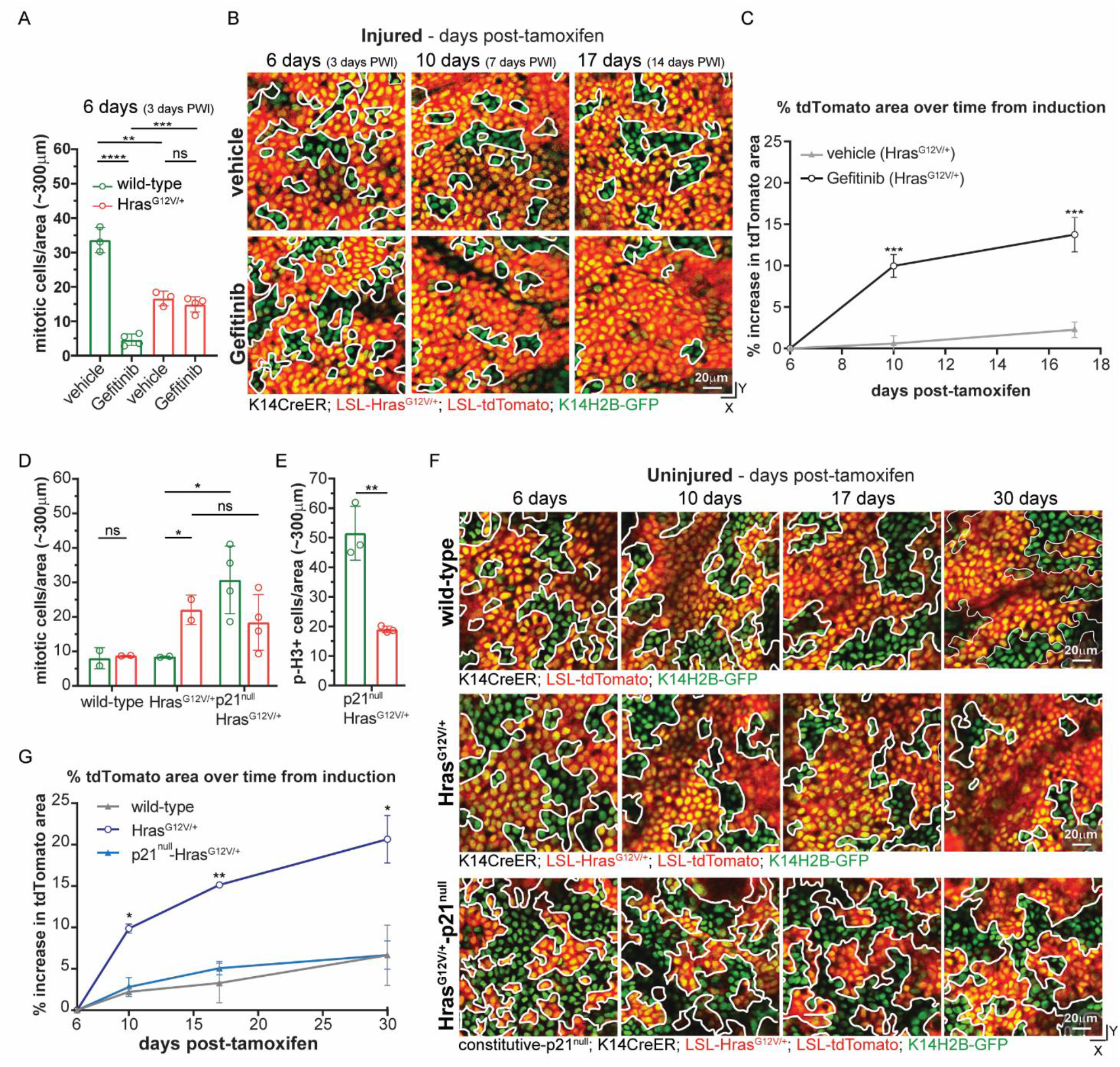
Increased wild-type cell proliferation is sufficient to suppresses the competitive advantage of Hras^G12V/+^ cells in mosaic skin. **A)** Quantification of mitotic figures in GFP+/tdTomato+ and GFP+/tdTomato− areas in both Hras^G12V/+^-mosaic treated daily with either vehicle or 200 mg/kg of Gefitinib (*see Materials and Methods*). The number of mitotic events was divided by the GFP+/tdTomato+ and GFP+/tdTomato− areas and multiplied by the total area analyzed (∼300 µm^2^) to compare the two populations. We analyzed three ∼300 µm^2^ areas for each mouse. n=3 vehicle and n=4 Gefitinib treated mice. **B)** Two-photon representative revisit images of the same area of the basal stem cell layer of the epidermis in injured Hras^G12V/+^-mosaic treated with vehicle (*top row*) or 200 mg/kg of Gefitinib (*bottom row*). White lines highlight the boundaries between GFP+/tdTomato+ and GFP+/tdTomato− populations. Scale bar=20 µm. **C)** Quantification of the percentage increase of tdTomato+ area at 6 days, 10 days and 17 days post-tamoxifen injection in injured Hras^G12V/+^-mosaic treated with vehicle or 200mg/kg of Gefitinib. Hras^G12V/+^ cells (GFP+/tdTomato+) reestablished their competitive advantage over wild-type neighbors (GFP+/tdTomato−) during injury-repair upon EGFR inhibition with Gefitinib. **D)** Quantification of mitotic figures in GFP+/tdTomato+ and GFP+/tdTomato− areas in wild-type-mosaic, Hras^G12V/+^-mosaic and constitutive-p21^null^-^HrasG12V/+^-mosaic. n=2 wild-type-mosaic and Hras^G12V/+^-mosaic mice (n=6 considering experiments in Fig. 2) and n=4 constitutive-p21^null^-Hras^G12V/+^-mosaic mice. **E)** Quantification of phospho-Histone-3 positive cells in GFP+/tdTomato+ and GFP+/tdTomato− areas in constitutive-p21^null^-Hras^G12V/+^-mosaic. The number of positive cells in the immunostaining was divided by the GFP+/tdTomato+ and GFP+/tdTomato− areas and multiplied by the total area analyzed (∼300 µm^2^) to compare the two populations. We analyzed three ∼300 µm^2^ areas for each mouse. n=3 mice for each condition. **F)** Two-photon representative revisit images of the same area of the basal stem cell layer of the epidermis in uninjured condition in wild-type-mosaic (*top row*), Hras^G12V/+^-mosaic (*middle row*) and constitutive-p21^null^-Hras^G12V/+^-mosaic (*bottom row*). White lines highlight the boundaries between GFP+/tdTomato+ and GFP+/tdTomato− populations. Scale bar=20 µm. **G)** Quantification of the percentage increase of tdTomato+ area at 6 days, 10 days, 17 days and 30 days post-tamoxifen injection in uninjured condition in wild-type-mosaic, Hras^G12V/+^-mosaic and constitutive-p21^null^-Hras^G12V/+^-mosaic. n=2 wild-type-mosaic mice and Hras^G12V/+^-mosaic mice (n=6 considering experiments in Fig. 2) and n=4 constitutive-p21^null^-Hras^G12V/+^-mosaic mice. Statistics: unpaired, ordinary one-way ANOVA between wild-type-mosaic, Hras^G12V/+^-mosaic and constitutive-p21^null^-Hras^G12V/+^-mosaic at different time points in uninjured condition. Unpaired, two-tailed *t-test* for comparison between different populations. *, P<0.05 and **. P<0.005. ns indicates not statistically significant. Data are represented as means and standard deviations.

Collectively, our data demonstrate that the EGFR/Ras signaling pathway is required to selectively increase wild-type cell proliferation after injury and to suppress Hras^G12V/+^ cell expansion in mosaic models.

### Loss of the cell cycle regulator p21 suppresses the competitive advantage of Hras^G12V/+^ cells in uninjured mosaic skin

To examine whether increased proliferation of wild-type cells is sufficient to suppress the competitive advantage of Ras-mutant cells in the absence of injury, we directly manipulated cell proliferation in the uninjured setting. The G1/S phase cyclin-dependent kinase inhibitor p21 is expressed in G1 phase cells to maintain skin epithelium homeostasis(*41, 42*), and its genetic ablation leads to increased proliferation in wild-type mouse skin epithelial cells *in vitro* and *in vivo*(*41-44*). Indeed, phospho-Histone-3 immunostaining of a p21^null^ model revealed substantially increased proliferation of basal stem cells, but not dermal cells, in uninjured mice (**Fig. S7C, D**). We hypothesized that Hras^G12V/+^ cells downregulate p21 expression to promote a high proliferative rate. Interestingly, p21 expression was significantly reduced at both mRNA and protein levels in Hras^G12V/+^-mosaic models compared to wild-type-mosaic models (**Fig. S8A, B, C, D, E**). Therefore, we reasoned that ubiquitous p21 deletion in the Hras^G12V/+^-mosaic model may selectively manipulate wild-type cells without affecting Hras^G12V/+^ cells, which express low levels of p21. To this end, we combined the p21^null^ model with the tamoxifen-inducible Hras^G12V/+^ model (*see Materials and Methods;* K14CreER; LSL-Hras^G12V/+^; LSL-tdTomato; constitutive p21^null^; K14H2B-GFP). Excitingly, p21 loss increased the proliferation of wild-type cells but not of Hras^G12V/+^ cells in p21^null^-Hras^G12V/+^-mosaic mice, mimicking the selective increase in wild-type cell proliferation during injury-repair of Hras^G12V/+^-mosaic mice (**Fig. 5D, E**). Importantly, the constitutive loss of p21 and increased proliferation were sufficient to suppress the competitive advantage of Hras^G12V/+^ cells over time in uninjured mice (**Fig. 5F, G, Fig. S7E)**, recapitulating the effects of injury-repair in Hras^G12V/+^-mosaic mice.

To explore the molecular mechanism that specifically increases the proliferation of wild-type cells but not Hras^G12V/+^ cells in the p21^null^ model and injury model, we probed the activation status of one of the key regulators of cell proliferation downstream of Ras, mitogen-activated protein kinase (MAPK, ERK1/2)(*45*). In uninjured skin, wild-type mice exhibited a lower level of activated ERK1/2 (phosphoERK1/2) when compared to the Hras^G12V/+^-mosaic and -max models, as expected (**Fig. S9A, B**). We observed increased levels of phosphoERK1/2 in wild-type mice that were undergoing injury-repair or that lacked p21 (**Fig. S9A, B**). However, the increase in phosphoERK1/2 was not significantly different in Hras^G12V/+^-mosaic and -max models in uninjured and injured conditions (**Fig. S9A, B**). Moreover, phosphoERK1/2 levels were similar after injury in all three models (wild-type, Hras^G12V/+^-mosaic models and Hras^G12V/+^-max, **Fig. S9A, B**). Lastly, the activation of Akt (phosphoAkt), another downstream target of the Ras pathway, was not significantly affected by either injury or p21 loss (**Fig. S9C**).

Overall, these data suggest that injury-repair or loss of p21 specifically increase the activity of a downstream effector of Ras, ERK1/2, in wild-type cells to increase their proliferation, enabling them to effectively suppress the competitive advantage of oncogenic Ras-mutant cells in mosaic mice.

## Discussion

Healthy tissues, including skin, harbor a number of somatic mutations, some of which are in known tumor driver genes(*1, 2, 46-48*). Models have shown that tumors can arise from the accumulation of multiple mutations, or from a lower mutational burden cooperating with additional exogenous insults, such as injury(*12-20, 25, 26*). We discovered that Ras-mutant cells break tissue architecture during injury-repair only when they represent nearly all the basal stem cells in the skin epidermis. In contrast, when Ras-mutant cells coexist with wild-type neighbors, injury selectively activates the endogenous proliferation program in wild-type cells only, which counteracts Ras-mutant cell expansion and oncogenesis (**Fig. 6**). Specifically, after an initial spike post-injury induction, the proliferation of wild-type cells equalizes to the level of Ras-mutant cells, higher than the proliferation of wild-type cells in homogeneous wild-type models. Neighboring wild-type cells exert a powerful defensive mechanism, even in the presence of a higher mutational burden from the most aggressive isoforms of the Ras family, Kras.

**Fig. 6:**
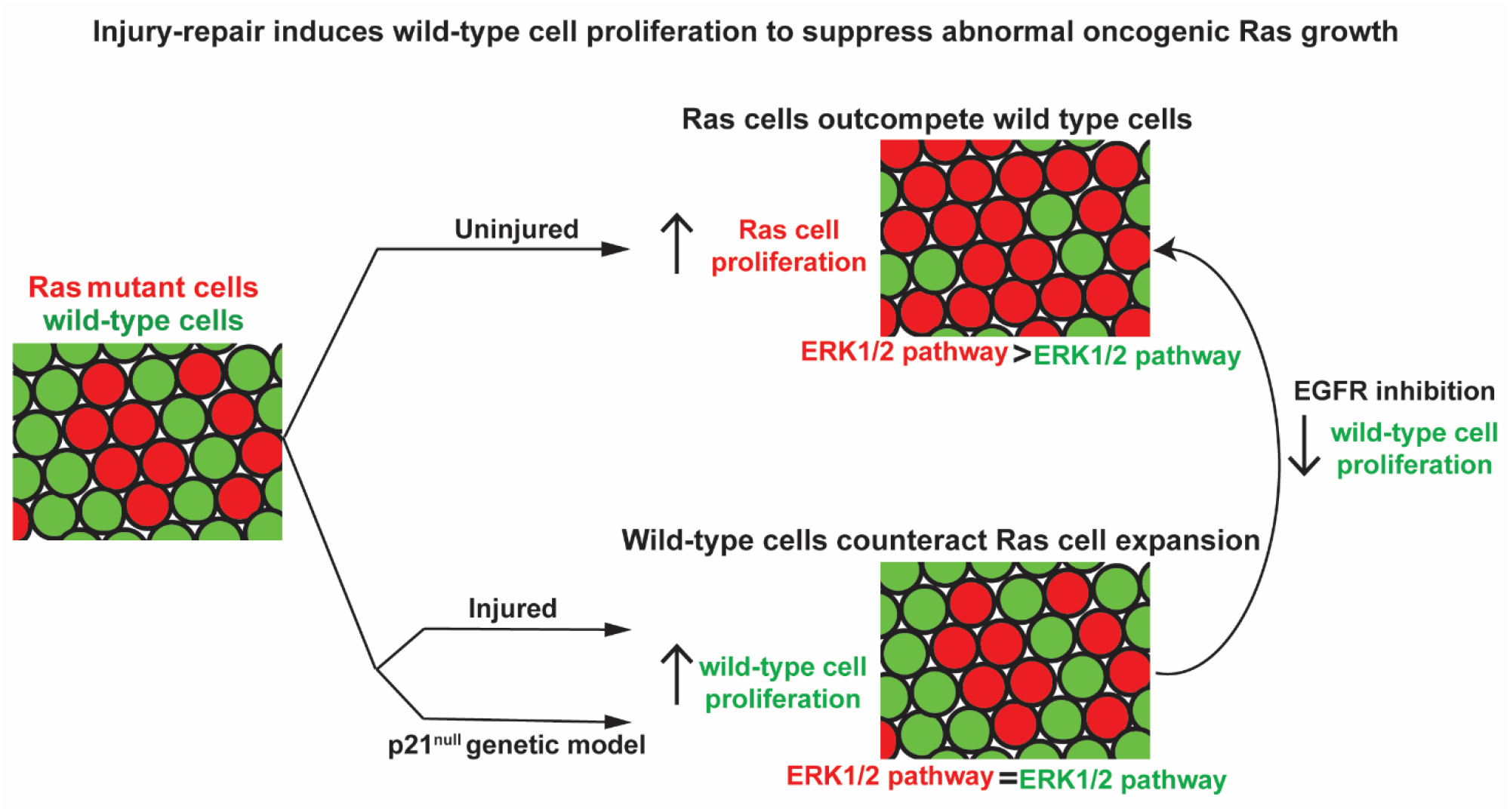
Injury-repair induces wild-type cell proliferation to suppress abnormal oncogenic Ras growth. In the uninjured mosaic skin epidermis, Ras epithelial cells integrate and expand, outcompeting wild-type neighbors. During injury-repair of mosaic skin, the competitive advantage of Ras cells over wild-type cells rather than enhanced is suppressed. Further, injury-induced Ras oncogenic growth do not develop. Injury induces wild-type cell proliferation, whereas Ras cell proliferation is unchanged. EGFR signaling pathway is crucial for increasing selectively wild-type cell proliferation to suppress Ras cell expansion during injury-repair. Ras cells are not sensitive to other pro-proliferative stimuli mediated by the EGFR pathway due to the constitutive activation of Ras, a downstream target of EGFR. Direct manipulation of proliferation, via constitutive p21 loss, revealed the sufficiency of wild-type cell proliferation in counteracting the competitive advantage of Ras cells even in the absence of the injury. Our data support a model whereby injury-repair and p21 loss increase the activity of ERK1/2, a Ras downstream pathway that controls cell proliferation. This leads to comparable ERK1/2 levels between wild-type and Ras models and results in an increase in wild-type cell proliferation that effectively suppresses the competitive advantage of Ras mutant cells.

We found that, although injury-repair is coordinated by various ligands/receptors, the EGFR/Ras signaling pathway emerges as key for selectively increasing wild-type cell proliferation to suppress Ras-mutant cell expansion. In the absence of injury, the constitutive loss-of-function of p21 also leads to a selective increase in proliferation of wild-type cells in uninjured mosaic skin, recapitulating the responses to injury and highlighting the sufficiency of wild-type cell proliferation as a protective mechanism against Ras-mutant cell expansion. Thus, genetic and environmental mechanisms enhance wild-type cell proliferation, whereas Ras-mutant cells are insensitive. Molecularly, our data suggest that the Hras^G12V^ mutation renders cells insensitive to pro-proliferative stimuli mediated by the EGFR/Ras pathway during injury-repair, in part because they already have high levels of phosphoERK1/2 activation, which promotes cell proliferation(*49*). Based on this observation, we propose that the fold change in Ras pathway activation before and after injury contribute to the selective capacity of wild-type cells to respond to pro-proliferative stimuli during injury-repair that are already maximized in Hras^G12V/+^ cells **(Fig. 6**).

Our findings have broad implications, given that Ras-mutant cells have competitive advantages over wild-type cells not only in uninjured skin, but also in other uninjured tissues, such as intestinal crypts and blood(*50-52*). Recent studies in a single layer epithelium *in vitro* and *in vivo* have shown that Ras^G12V^-mutated cells are apically extruded when surrounded by wild-type epithelial cells(*53-55*). In the stratified epithelium of skin epidermis, our studies and others showed that Ras-mutant cells expand to outcompete wild-type neighbors and integrate in a healthy tissue, suggesting a different mode of cell competition compared to the systems above(*9-11*). We reconcile the evidence above by noting that different tissues preserve and maintain their specific architecture through distinct cell behaviors. Endogenous behaviors for monolayer epithelia are proliferation and extrusion, whereas multilayer epithelia rely on proliferation and delamination/differentiation. Thus, in a stratified epithelium, differentiation is analogous to extrusion in monolayer epithelia. Consistent with this reasoning, our evidence suggests that apoptosis, an ectopic behavior in the adult skin epithelium, is not involved in the competition between wild-type and Hras^G12V/+^ cells.

Traditional therapeutic approaches used for cancer treatment involve suppressing the proliferation of all cells, both mutant and wild-type cells. While these approaches restrain tumor expansion, they also impair the opportunity for the tissue to deploy natural defenses, such as selective promotion of wild-type cell proliferation. The next step towards an effective therapeutic treatment would be to determine how to promote the proliferative advantage bestowed on wild-type cells in the injury environment or in the pro-proliferative p21^null^ state. Our data would argue that in precancerous states, EGFR activation, such as through EGF treatment, might provide a competitive advantage to wild-type cells in the presence of neighbors expressing the constitutively active form of Ras oncogene. Collectively, this work provides way forward for future research and clinical application to shift the focus on the mechanism that empower wild-type cells in the competition with their mutated opponent.

## Acknowledgements

We thank all members of the Greco lab, Beronja S (Fred Hutchinson Cancer Research Center, Seattle, WA), Regot S (Johns Hopkins University School of Medicine, Baltimore, MD), Politi K and Hu B (Yale university, New Haven, CT) and Andersen A (Life science editor) for critical feedback on the manuscript. We thank Muzumdar MD (Yale University, New Haven, CT) for the LSL-Kras^G12D/+^ mice.

## Author contributions

SG and VG designed experiments and wrote the manuscript. SG performed two-photon imaging, epidermal preparation staining, mouse genetics, image analyses, designed the scRNA-seq experiment and assisted with analyses. DG assisted with IMARIS and MatLab analyses. NR and KA assisted with data analyses of scRNA-seq data. MK assisted with interpretation of scRNA-seq data. SY assisted with IMARIS data analysis. CM assisted with epidermal preparation and staining. EL and TX assisted with whole mount tissue and OCT staining. KS assisted with clinical diagnosis of histological images and provided critical feedback on the manuscript.

## Data and materials availability

All data from this study are available from the authors on request. scRNA-seq data will be available in Gene Expression Omnibus (GEO) in the accepted manuscript.

## Supplementary Materials

Materials and Methods

Figs. S1 to S9

References (*20, 21, 23, 24, 56-69*)

Movies S1 to S4

